# The remarkable robustness of surrogate gradient learning for instilling complex function in spiking neural networks

**DOI:** 10.1101/2020.06.29.176925

**Authors:** Friedemann Zenke, Tim P. Vogels

## Abstract

Brains process information in spiking neural networks. Their intricate connections shape the diverse functions these networks perform. In comparison, the functional capabilities of models of spiking networks are still rudimentary. This shortcoming is mainly due to the lack of insight and practical algorithms to construct the necessary connectivity. Any such algorithm typically attempts to build networks by iteratively reducing the error compared to a desired output. But assigning credit to hidden units in multi-layered spiking networks has remained challenging due to the non-differentiable nonlinearity of spikes. To avoid this issue, one can employ surrogate gradients to discover the required connectivity in spiking network models. However, the choice of a surrogate is not unique, raising the question of how its implementation influences the effectiveness of the method. Here, we use numerical simulations to systematically study how essential design parameters of surrogate gradients impact learning performance on a range of classification problems. We show that surrogate gradient learning is robust to different shapes of underlying surrogate derivatives, but the choice of the derivative’s scale can substantially affect learning performance. When we combine surrogate gradients with a suitable activity regularization technique, robust information processing can be achieved in spiking networks even at the sparse activity limit. Our study provides a systematic account of the remarkable robustness of surrogate gradient learning and serves as a practical guide to model functional spiking neural networks.

## Introduction

The computational power of deep neural networks [1, 2] has reinvigorated interest in using in-silico systems to study information processing in the brain [3, 4]. For instance, performance-optimized artificial neural networks bear striking representational similarity with the visual system [5–11] and serve to formulate hypotheses about their mechanistic underpinnings. Similarly, the activity of artificial recurrent neural networks optimized to solve cognitive tasks resembles cortical activity in prefrontal [12, 13], medial frontal [14], and motor areas [15, 16] and inspire comparison and lively discussion.

All of these studies rely on conventional artificial neural networks with graded activation functions as commonly used in machine learning. The recipe for building a deep neural network is straight-forward. The value of a scalar loss function defined at the output of the network is decreased through gradient descent. They differ from biological neural networks in important respects. For instance, they lack cell type diversity and do not obey Dale’s law while ignoring the fact that the brain uses spiking neurons. We generally accept these flaws because we do not know how to construct more complicated networks. For instance, gradient descent only works when the involved system is differentiable. This is not the case for spiking neural networks (SNNs).

Surrogate gradients have emerged as a solution to build functional SNNs capable of solving complex information processing problems [17–24]. To that end, the actual derivative of a spike, which appears in the analytic expressions of the gradients, is replaced by any well-behaved function. There are many possible choices of such surrogate derivatives, and consequently, the resulting surrogate gradient is, unlike the true gradient of a system, not unique. A number of studies have successfully applied different instances of surrogate derivatives to various problem sets [19, 21, 22, 25–27]. While this suggests that the method does not crucially depend on the specific choice of surrogate derivative, we know relatively little about how the choice of surrogate gradient impacts the effectiveness and whether some choices are better than others. Previous studies did not address this question because they solved different computational problems, thus precluding a direct comparison. In this article, we address this issue by providing benchmarks for comparing the trainability of SNNs on a range of supervised learning tasks and systematically vary the shape and scale of the surrogate derivative used for training networks on the *same* task.

## Results

To systematically evaluate the performance of surrogate gradients, we sought to repeatedly train the same network on the same problem while changing the surrogate gradient. Towards this end, we required a demanding spike-based classification problem with a small computational footprint to serve as benchmark. There are only few established benchmarks for SNNs. One approach is to use analog-valued machine learning datasets as input currents directly [17] or to first convert them to Poisson input spike trains [18, 20]. These input paradigms, however, do not fully capitalize on the ability to encode information in spike timing, an important aspect of spiking processing. Gütig [28] addressed this point with the Tempotron by classifying randomly generated spike timing patterns in which each input neuron fires a single spike. Yet, completely random timing, precludes the possibility to assess generalization performance, i.e., the ability to generalize to previously unseen inputs. To assess if SNNs could learn to categorize spike patterns and generalize to unseed patterns, we created a number of synthetic classification datasets with added temporal structure. Specifically, we created spike rasters for a given set of input afferents. Each afferent only fired one spike, and the spike times of all afferents were constrained to lie on a low dimensional smooth random manifold in the space of all possible spike timings. All data points from the same manifold were defined as part of the same input class, whereas different manifolds correspond to different classes.

The spike-timing manifold approach has several advantages: First, the temporal structure in the data permits studying generalization, a decisive advantage over using purely random spike patterns. Second, the task complexity is seamlessly adjustable by tuning the number of afferents, manifold’s smoothness parameter *α* (Fig. 1a), the intrinsic manifold dimension *D*, and the number of classes *n* (Fig. 1b). Third, we ensure that each input neuron spikes exactly once (Fig. 1c), guaranteeing that, the resulting datasets are purely spike-timing-dependent and thus cannot be classified from firing rate information. Finally, sampling an arbitrary number of data points from each class is computationally cheap and it is equally easy to generate an arbitrary number of different datasets with comparable properties.

**Fig 1.**
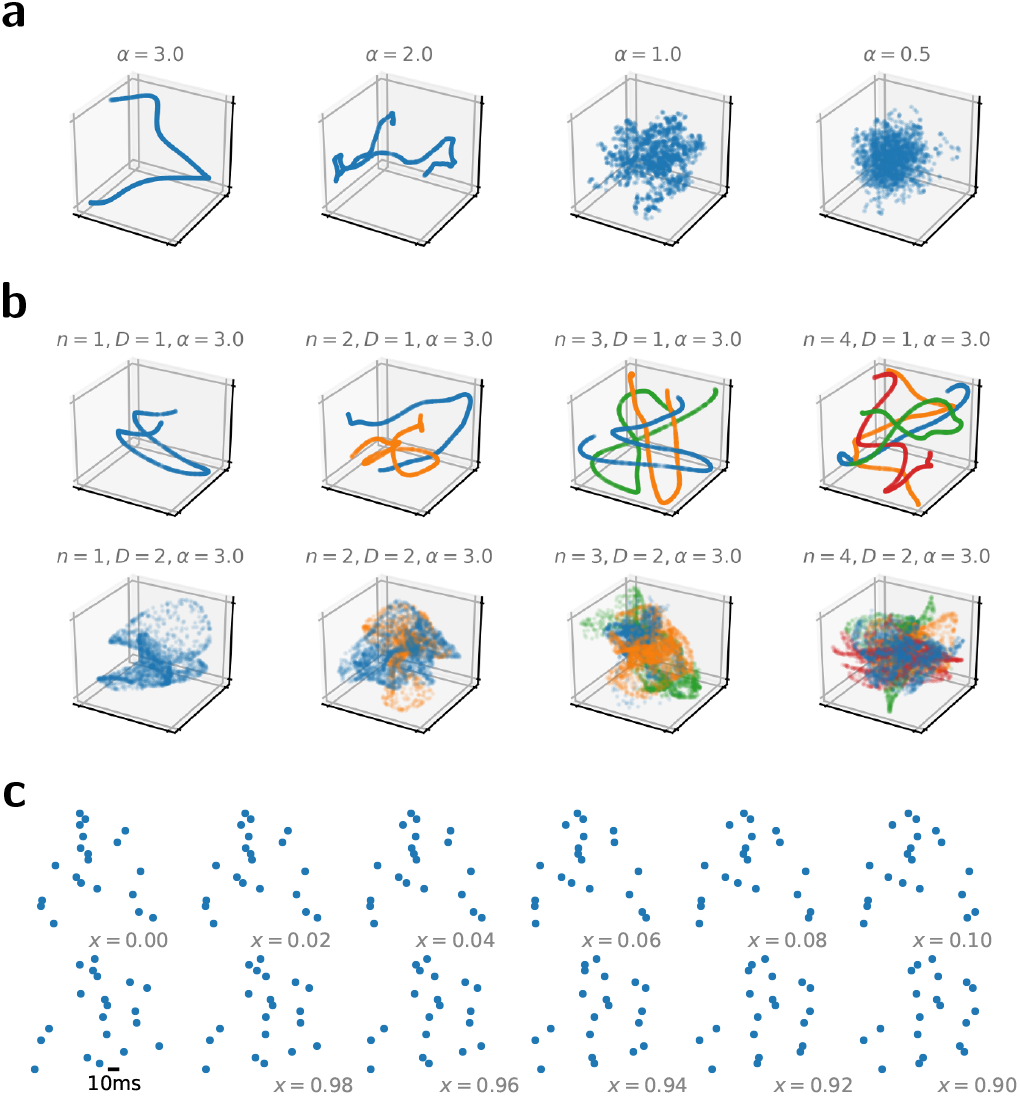
Smooth random manifolds provide a flexible way of generating synthetic spike-timing datasets. **(a)** Four one-dimensional example manifolds for different smoothness parameters *α* in a three-dimensional embedding space. From each manifold, we plotted 1000 random data points. **(b)** Same as in (a), but keeping *α* = 3 fixed while changing the manifold-dimension *D* and the number of random manifolds (different colors). By sampling different random manifolds, it is straight forward to build synthetic multi-way classification tasks. **(c)** Spike raster plots corresponding to twelve samples along the intrinsic manifold coordinate *x* of a one-dimensional smooth random manifold (*α* = 3) whereby we interpreted the embedding space coordinates as firing times of the individual neurons.

To demonstrate the validity of our approach, we tested it on a SNN with a single hidden layer on a simple two-way classification problem (Fig. 2a; Methods). We modeled the units of the hidden layer as current-based leaky integrate-and-fire neurons. Between layers, the connectivity was strictly feed-forward and all-to-all. The output layer consisted of two leaky integrators that did not spike, allowing us to compute the maximum of the membrane potential [29] and to interpret these values as the inputs for a standard classification loss function for supervised learning (Methods). In this setup, the readout unit with the highest activity level signals the putative class-membership of each input [23].

**Fig 2.**
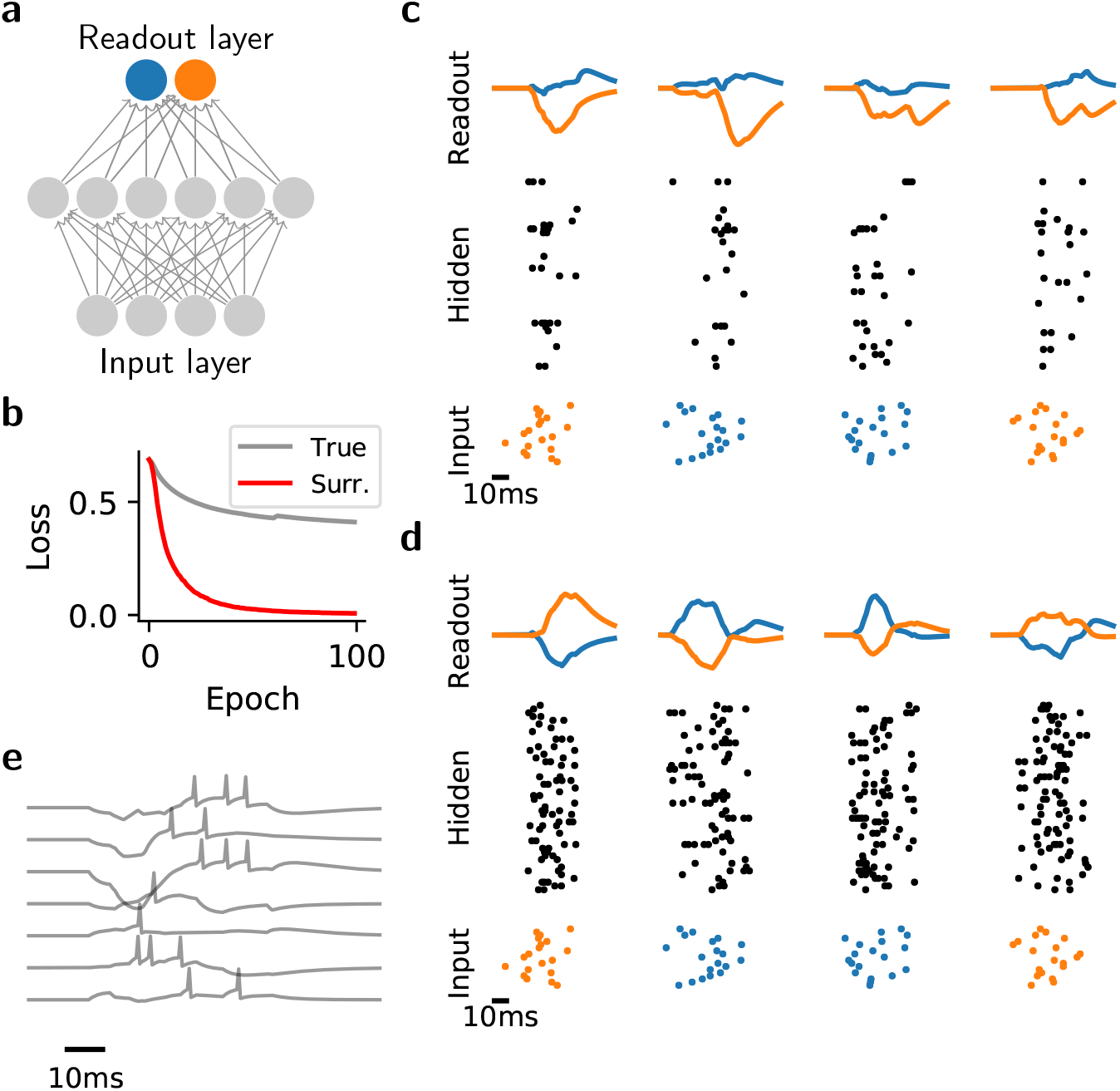
Surrogate gradient descent allows building functional SNNs. **(a)** Sketch of the network model with two readout units at the top. **(b)** Learning curves of the network when using the actual gradient (“true”, gray) or a surrogate gradient (red) to train an SNN on a binary random manifold classification problem. **(c)** Snapshot of network activity before training. Bottom: Spike raster of the input layer activity. Four different inputs corresponding to two different classes are plotted in time (orange/blue). Middle: Spike raster of the hidden layer activity. Top: Readout unit membrane potential. The network erroneously classifies the two “orange” inputs as belonging to the “blue” class, as can be read off from the maximum activity of its readout units. **(d)** Same as in c, but following training of the network with surrogate gradient descent. **(e)** Example membrane potential traces from 7 randomly selected hidden layer neurons during a single trial.

We first confirmed that learning is poor when we used the actual gradient. To that end, we computed it using the derivative of the hard threshold nonlinearity of the spikes. As expected, the hard threshold nonlinearity prevented gradient flow into the hidden layer [24], and consequently lead to poor performance (Fig. 2b,c). In contrast, when we used surrogate gradients to train the same network, the problem disappeared. Learning took place in both hidden and output layer and resulted in a substantial reduction of the loss function (Fig. 2b–e).

### Surrogate gradient learning is robust to the shape of the surrogate derivative

A necessary ingredient of surrogate grading learning is a suitable surrogate derivative. To study the effect of the surrogate derivative comparably, we generated a random manifold dataset with ten classes. We chose the remaining parameters, i.e., the number of input units, the manifold dimension, and the smoothness *α*, to make the problem impossible to solve for a network without hidden layer while at the same time keeping the computational burden minimal. We trained multiple instances of the same network with a single hidden layer (*n*_h_ = 1) using the derivative of a fast sigmoid as a surrogate derivative (Fig. 3a “SuperSpike”; [25]) on this dataset. In each run, we kept both the dataset and the initial parameters of the model fixed, but varied the slope parameter *β* of the surrogate. For each value of *β* we performed a parameter sweep over the learning rate *η*. Following training, we measured the classification accuracy on held-out data. This search revealed an extensive parameter regime of *β* and *η* in which the system was able to solve the problem with high accuracy (Fig. 3b). The addition of a second hidden layer only marginally improved upon this result, whereas, as expected, a network without hidden layer performed poorly (Fig. 3c). The extent of the parameter regime yielding high performance suggests remarkable robustness of surrogate gradient learning to changes in the “steepness” of the surrogate derivative. While a steep approach to the threshold could be seen as a closer, hence better approximation of the actual derivative of the spike, the surrogate gradient remains largely unaffected by how *closely* the function resembles the exact derivative as long as it is not a constant.

**Fig 3.**
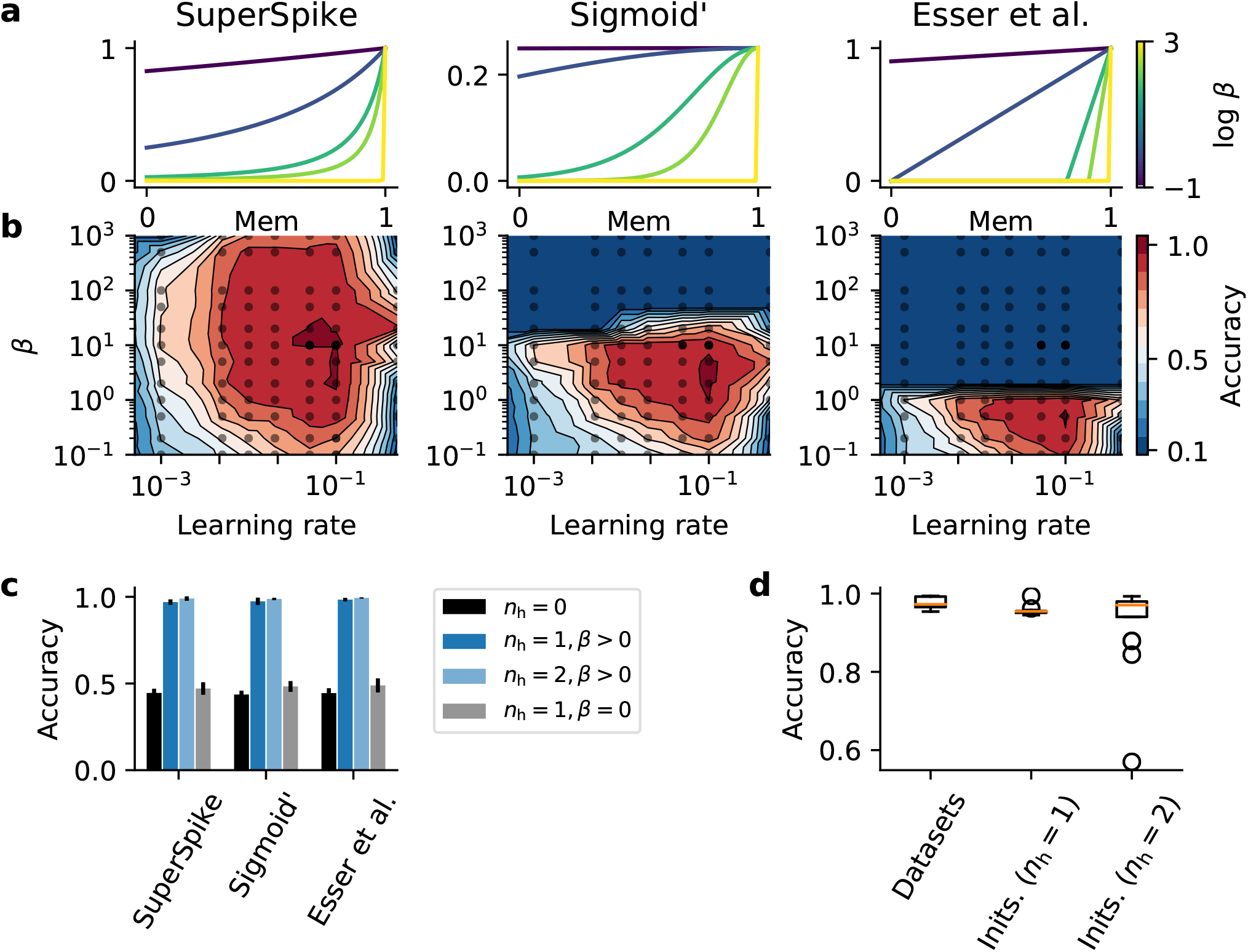
Surrogate gradient learning is robust to the shape of the surrogate derivative. **(a)** Three different surrogate derivative shapes that have been used for training on the synthetic smooth random manifold spiking dataset. From left to right: SuperSpike [25], the derivative of a fast sigmoid function, Sigma′, the derivative of a standard sigmoid function, and “Esser et al.”, a piece-wise linear function [19, 22]. Colors correspond to different values of the slope parameter *β*. **(b)** Accuracy on held-out data as a function of the learning rate *η* and the slope *β* for the corresponding surrogates in (a) for a network with one hidden layer (*n_h_* = 1). **(c)** Test accuracy for the five best parameter combinations obtained from a grid search, as shown in (b) for different surrogates and numbers of hidden layers *n_h_*. While a network without hidden layer is unable to solve the classification problem (black), networks trained with a wide range of different surrogates and slope parameters (*β* > 0) have no problem solving the task with high accuracy (shades of blue). However, the problem is not solved with high accuracy by a network with a hidden layer in which the surrogate derivative was a constant during training (*β* = 0; gray). Error bars correspond to the standard deviation (*n* = 5). **(d)** Whisker plot of classification accuracy for a network with one hidden layer over five different realizations of the random manifold datasets (“Datasets”) and for the same dataset, but using different weight initializations (“Inits.”) in networks with either one (*n*_h_ = 1) or two (*n*_h_ = 2) hidden layers.

Next we tested different surrogate derivative shapes, namely a standard sigmoid (Sigmoid′) and piece-wise linear function (Esser et al.; Fig. 3a; [19, 22]). This manipulation led to a reduction of the size of the parameter regime in *β* in which the network was able to perform the task, which is presumably due to vanishing gradients [30]. However, there was no substantial reduction in maximum performance (Fig. 3b,c). Using a piece-wise linear surrogate derivative (Esser et al.) led to a further reduction of viable parameters *β* (Fig. 3b), but did not affect maximum performance, regardless of whether we used one or two hidden layers (Fig. 3c). To check whether a surrogate derivative was required at all for solving the random manifold problem, we assayed the learning performance for *β* = 0, which corresponds to setting the function to 1. This change resulted in a significant drop in performance comparable to a network without hidden units (Fig. 3c) suggesting that a nonlinear voltage-dependence is crucial to learn useful hidden layer representations. Finally, we confirmed that these findings were robust to different initial network parameters and datasets (Fig. 3d) apart from only a few outliers with low performance in the two hidden layer case (*n_h_* = 2). These outliers point at the vital role of proper initialization [31, 32].

### Surrogate gradient learning in recurrent networks is sensitive to the scale of the surrogate derivative

In most studies that rely on surrogate gradients, the surrogate derivative is normalized to 1 [19, 21, 22, 24, 25] (cf. Fig. 3a), markedly different from the actual derivative of a spiking threshold which is infinite (Fig. 5a). The scale of the substituted derivative has a strong effect on the surrogate gradient due to both explicit and implicit forms of recurrence in SNNs (Fig. 4; [24]). Most notably, the scale may determine whether gradients vanish or explode [30]. However, it is unclear to what extent such differences translate into performance changes of the trained networks.

**Fig 4.**
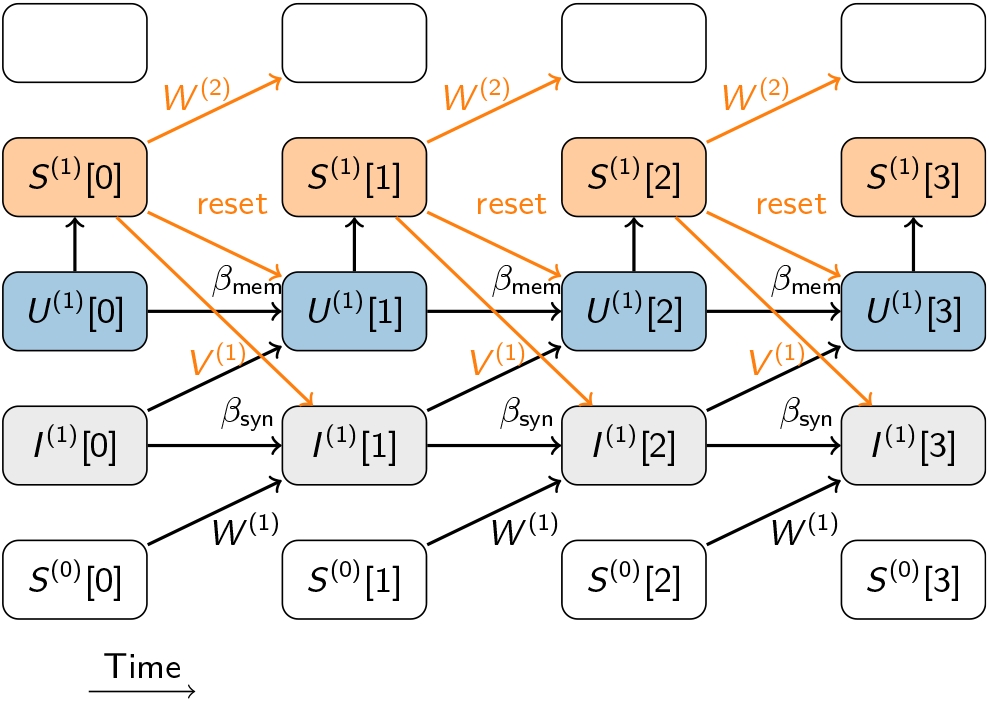
SNNs can be have both implicit and explicit recurrence. Schematic of the computational graph of a single SNN layer composed of leaky integrate-and-fire (LIF) neurons (see Methods). Input spike trains *S*^(0)^ enter at the bottom and affect the synaptic current variable *I*^(1)^ through the feed-forward weights *W* ^(1)^. Time flows from left to right. Any link that connects temporally adjacent nodes in the graph constitute a form of recurrence in the computation whereby the synaptic connections *V* ^(1)^ contribute explicit recurrence to the graph. Implicit recurrence is contributed, for instance, by the decay of synaptic current variables and the membrane potentials *U* ^(1)^. Additionally, the spike reset contributes another form of implicit recurrence by coupling the future states to the output spike train *S*^(1)^. Recurrences involving the surrogate derivative, e.g. the reset, depend on both the shape and the scale of the surrogate chosen and can substantially alter the surrogate gradient.

**Fig 5.**
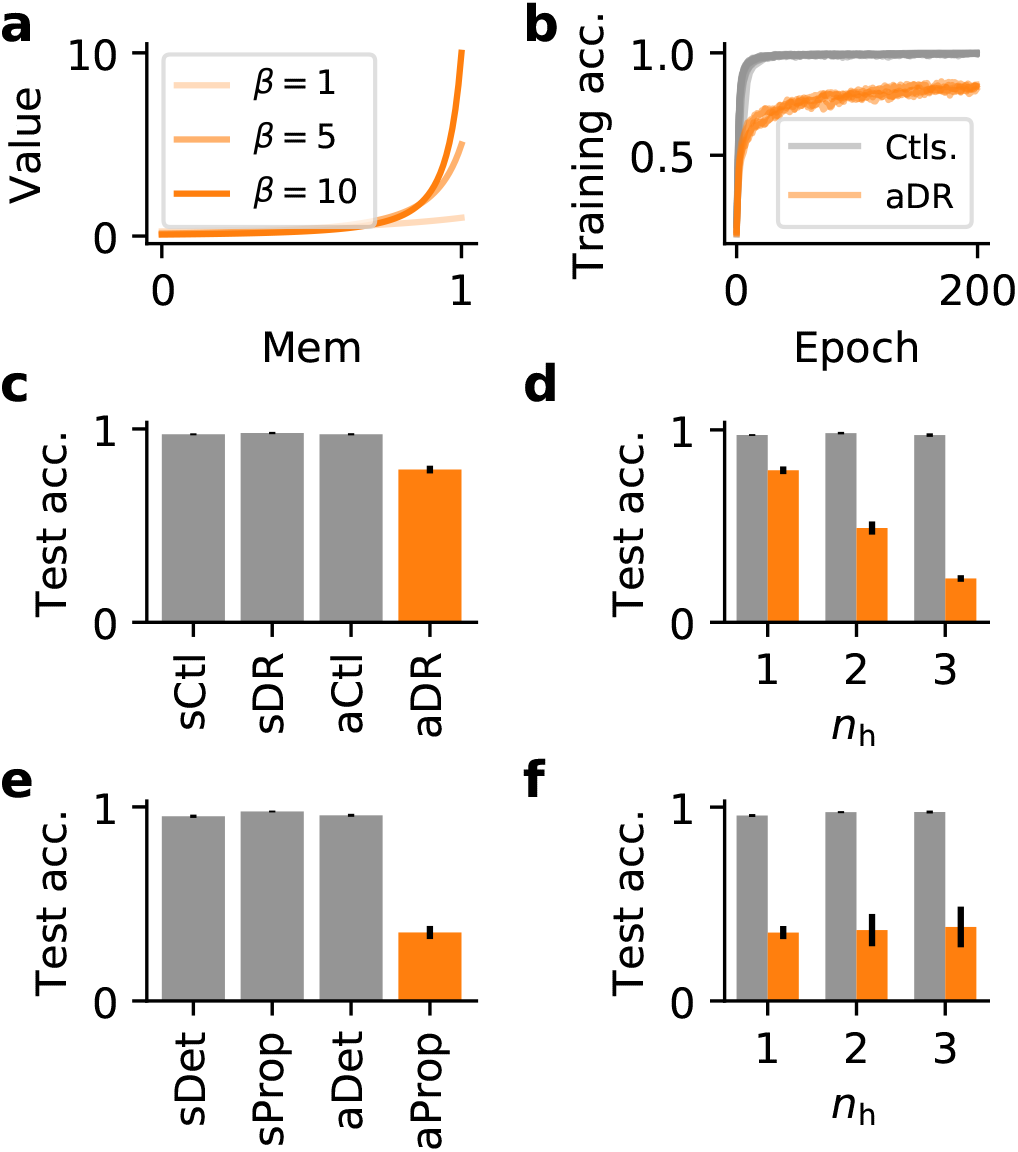
Surrogate gradient learning is sensitive to the scale of the surrogate derivative. **(a)** Illustration of pseudo derivatives *σ′* that converge toward the actual derivative of a hard spike threshold *β* → ∞. Note that in contrast to Fig. 3a, their maximum value grows as *β* increases. **(b)** Training accuracy of several spiking networks (*n_h_* = 1) during training on a synthetic classification task. The gray curves comprise control networks in which the surrogate derivative was either normalized to one or in which we used an asymptotic surrogate derivative, but prevented surrogate gradients from flowing through the spike reset. Orange curves correspond to networks with asymptotic pseudo derivatives with differentiable spike reset (aDR). In all cases, we plot the five best performing learning curves obtained from an extensive grid-search over *β* and the learning rate *η* (cf. Fig. 3). **(c)** Quantification of the test accuracy of the different learning curves shown in (b). We trained all networks using a SuperSpike nonlinearity. The reset term was either ignored (sCtl), or in which a differentiable reset was used (vDR). Similarly, we considered an asymptotic variant of SuperSpike that does converge toward the exact derivative of a step function for *β* → ∞, without (aCtl) or with a differentiable reset term (aDR). The results shown correspond to the ten best results from a grid search. The error bars denote the standard deviation. **(d)** A similar comparison of control cases in which reset terms were ignored (gray) or could contribute to the surrogate gradient (orange) for different numbers of hidden layers. **(e)** Test accuracy as in (c), but comparing SuperSpike (v) and the asymptotic (a) case in which gradients can flow through recurrent connections (Prop) versus the detached case (Ctl). **(f)** Test accuracy for asymptotic SuperSpike as a function of the number of hidden layers for networks in which gradients were flowing through recurrent connections (orange) versus the detached case (gray).

To gain a better understanding of how derivative scales larger than 1 affect surrogate gradient learning, we trained networks on a specific random manifold task, an *asymptotic* version of the SuperSpike surrogate (aCtl; Fig. 3) and our well-tested standard SuperSpike function (sCtl; Fig. 3) as a control (sCtl). As we expected the difference in scale to manifest itself primarily in the presence of recurrence, we compared networks in which we treated the spike reset as differentiable (DR) with networks in which its contribution was ignored by *detaching* it from the computational graph. Technically speaking, we prevented PyTorch’s auto-differentiation routines [33] from considering connections in the computational graph that correspond to the spike reset when computing the gradient with back-propagation through time (BPTT) (see Methods Eq. (1)). While both the normalized (sCtl) as well as the asymptotic surrogate (aCtl), performed equally well when the reset was detached, combining a differentiable reset with an asymptotic surrogate derivative (aDR) lead to impaired performance (Fig. 5b,c). This adverse effect on learning was amplified in deeper networks (Fig. 5d). Thus, the scale of the surrogate derivative plays indeed an important role for learning success if implicit recurrence, as contributed by the spike reset, is present in the network.

Since the spike reset constitutes a specific form of implicit recurrence (cf. Fig. 4), we were wondering whether we would observe a similar phenomenon for explicit recurrence through recurrent synaptic connections. To that end, we repeated the above performance measurements in networks with recurrent connections, but kept the spike reset term detached to prevent gradient flow. We observed a small but measurable reduction in accuracy for the best performing networks (Fig. 5c,e). Importantly, however, there was a substantial decrease in classification performance when gradients were allowed to flow through recurrent connections, *and* the asymptotic SuperSpike variant was used (Fig. 5e). Unlike in the differentiable reset case, the effect was severe enough to drop network performance to chance level even for a network with a single hidden layer (Fig. 5f). In summary, surrogate gradients are sensitive to the scale of the surrogate derivative. More specifically, when the scale of the surrogate derivative is too large, while either implicit or explicit recurrence are present in the network, the effect on learning can be detrimental.

### Surrogate gradient learning is robust to changes in the loss functions, input paradigms, and datasets

So far we investigated synthetic random manifold datasets in strictly feed-forward networks trained with loss functions that were defined on the maximum over time (Max) of the readout units. Next, we performed additional simulations in which the loss was computed by summation over time (“Sum”; see Methods). Based on our findings above, we limited our analysis to the SuperSpike surrogate with *β* = 10 and detached reset terms. As before, we performed a grid search over the learning rate *η* and selected the ten best performing models using held-out validation data. We then computed their classification performance on a separate test set. We repeated our simulation experiments on the above random manifold task and did not observe any substantial difference in the accuracy for the Max and Sum type readout heads (Fig. 6a).

**Fig 6.**
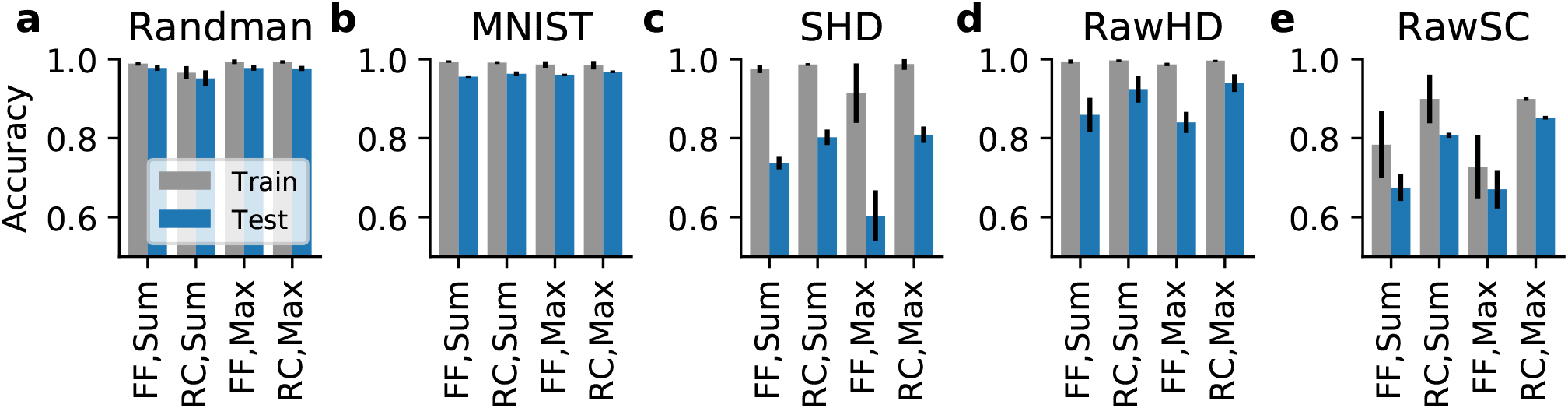
Surrogate gradient learning is effective on different loss functions, input paradigms, and datasets. Bar plots showing the test classification accuracy for different datasets (a–e). The plots further distinguish models by their readout configuration (Sum vs. Max) and whether they use purely feed-forward (FF) or explicitly recurrent (RC) synaptic connectivity. Each bar corresponds to the mean over the ten best performing models on held-out validation data and error bars signify the standard deviation.

To check the validity of these findings on a different dataset, we trained networks on MNIST by converting pixel values into spike latencies (Fig. 7a,b; Methods). In this paradigm, each input neuron fires either a single or no spike for each input. The networks reached a test accuracy of (97.4 ± 0.2) %, which is comparable to a conventional artificial neural network with the same number of neurons and hidden layers and to previous SNN studies using temporal coding [34]. We did not observe any discernible performance differences between the two readout types we tested (Fig. 6b).

**Fig 7.**
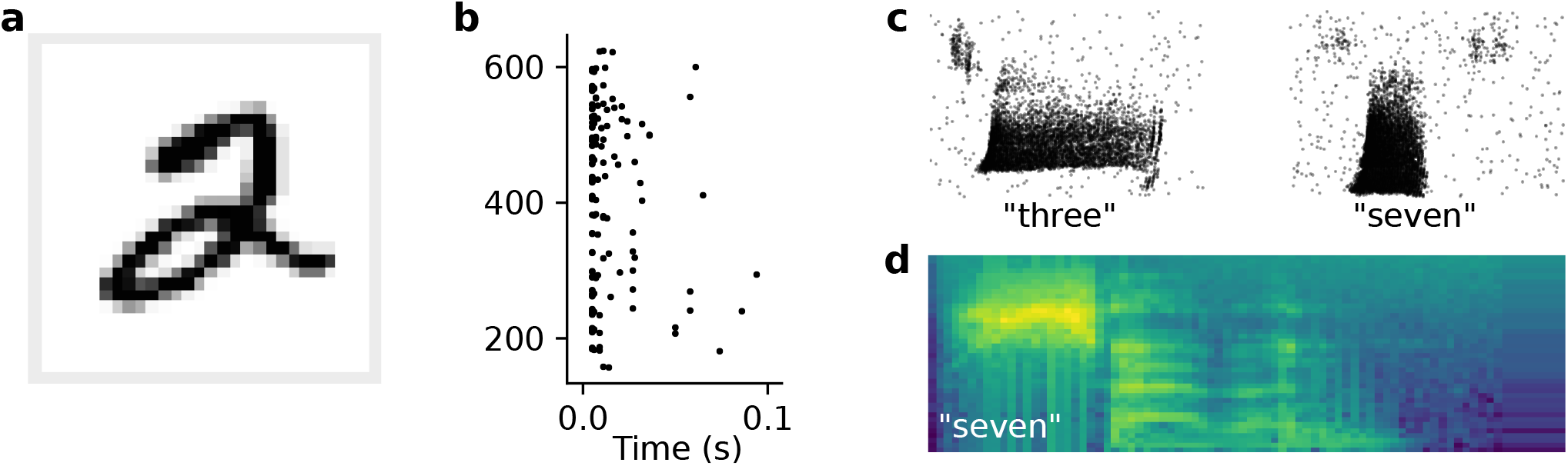
Examples of different input paradigms. **(a)** One example image from MNIST hand-written digit dataset. Spike raster plot of the corresponding spike latency encoding for the 28 × 28 = 784 input neurons. **(c)** Spike raster of two example inputs for the spoken digits “three” and “seven” taken from the SHD dataset [23]. **(d)** Mel-scaled spectrogram of an utterance of the number “seven” as used for simulations using raw audio input.

Further, to study the effect of explicit recurrence, we ran separate experiments for recurrently connected networks (RC). Importantly, we did not observe any substantial performance differences between strictly feed-forward and recurrently connected networks either (Fig. 6a,b).

We speculated that the absence of an effect may be due to the short duration of the input spike trains considered so far (~ 50ms). Recurrent connections are typically thought to provide neural networks with longer timescale dynamics, effectively giving the network a working memory. Thus, the beneficial effects of recurrent connections may only emerge when using stimuli of longer duration, with potentially multiple spikes from any given input neuron.

To test this hypothesis, we trained networks on the Spiking Heidelberg Digits (SHD) dataset [23], which consists of simulated input spikes from the auditory pathway of varied duration between 0.6–1.4s (Fig. 7c). Indeed, we found that the best performing models in this case were recurrent and achieving state-of-the-art classification accuracy of (0.82 ± 0.02) % (Fig. 6c). These data are consistent with the notion that working memory plays a vital role for the classification of longer input patterns.

### Surrogate gradient learning in networks with current-based input

Until now, we considered spike-based datasets. While spiking inputs are arguably the most natural input to SNNs, they come with an important caveat. All spiking datasets assume a specific encoding model, which is used to convert analog input data into a spiking representation. The chosen model, however, may not be optimal and thus adversely affect classification performance. To avoid this issue, we sought to *learn* the spike encoding by directly feeding current-based input to a set of spiking units [35]. To test this idea, we converted the raw audio data of the Heidelberg Digits [23] to Mel-spaced spectrograms (7d; Methods). To reduce overfitting, we decreased the number of channels and time steps to values commonly used in artificial speech recognition systems. Specifically, we used 40 channels and 80 time frames, corresponding to compression in time by a factor of about five (Methods). The networks trained on this RawHD dataset showed reduced overfitting, yet still benefited from recurrent connections compared to strictly feed-forward networks (Fig. 6d). Concretely, recurrent networks reached (94 ± 2) % test accuracy, whereas the feed-forward networks reached only (84 ± 3) %. In agreement with the results on Randman and MNIST, there were no significant differences between the Sum and Max readout configurations (Fig. 6).

We wanted to know whether the discrepancy between feed-forward and recurrent networks would increase for more challenging data sets. This was made possible by the performance gain from reducing the input dimension to 40 channels, which allowed us to train SNNs on the larger Speech Command dataset ([36]; Methods). This dataset comprises over 100k utterances from 35 classes, including “yes”, “no”, and “left”. In contrast to the original intended use for keyword spotting, here we assayed its top-1 classification accuracy over all classes, a more challenging problem than accurate detection of only a subset of words. The best SNNs achieved (85.3 ± 0.3) % on this challenging benchmark (Fig. 6e). On the same task, the spiking feed-forward network performed at (70 ± 2) %. There was a clear benefit from adding recurrent connections to the network, with the performance of the Max ((85.3 ± 0.3) %) being slightly better than the Sum ((80.7 ± 0.4) %) readout configuration.

These findings illustrate that surrogate gradient learning is robust to changes in the input paradigm, including spiking and non-spiking datasets. For more complex datasets, recurrently connected networks performed better than strictly feed-forward networks. Finally, in the majority of cases, surrogate gradient learning was robust to the details of how the output loss was defined.

### Optimal sparse spiking activity levels in SNNs

Up to now, we have focused on maximizing classification accuracy while ignoring the emerging activity levels in the resulting SNNs. However, we found that for some solutions the neurons in these networks displayed implausibly high firing rates (Fig. 8a). Experimental results suggest that most biological networks exhibit sparse spiking activity, a feature that is presumed to underlie their superior energy-efficiency [37–42].

**Fig 8.**
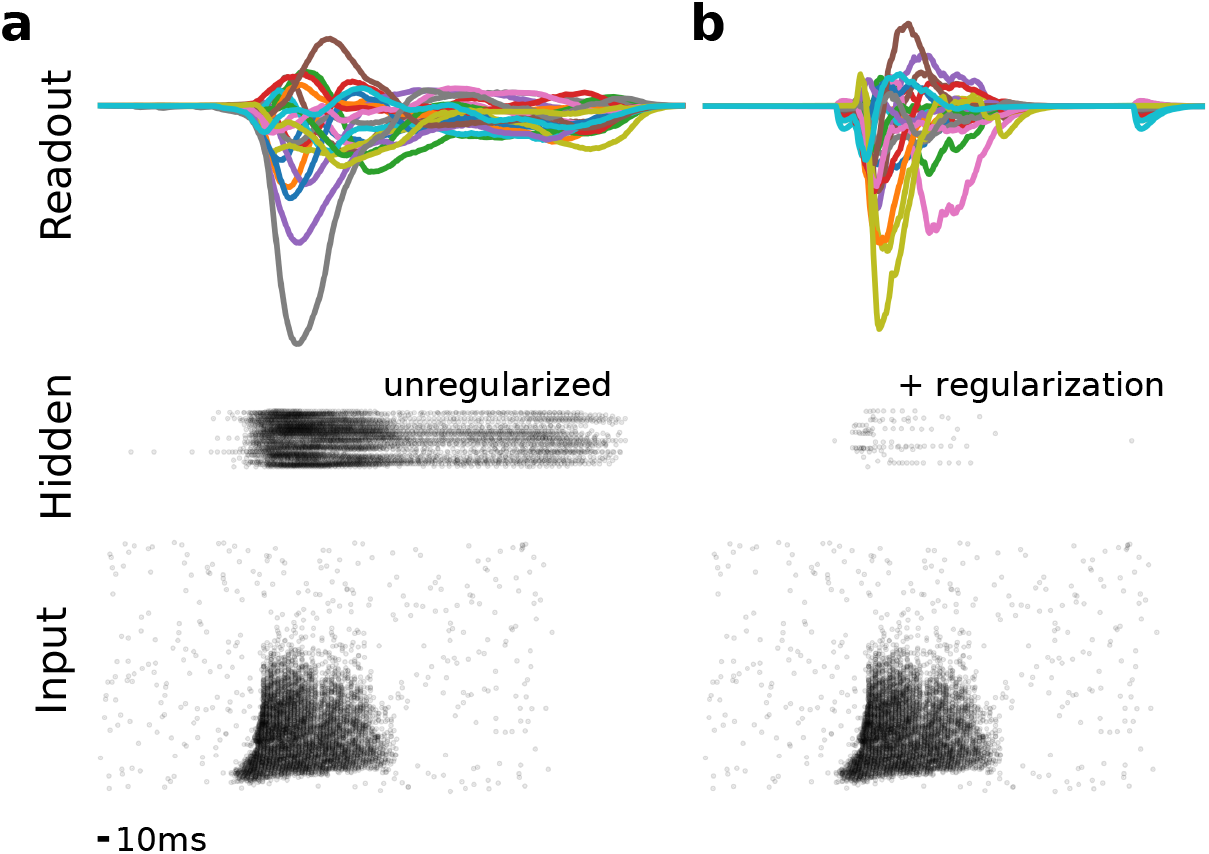
Activity regularization renders hidden layer activity sparse while maintaining functionality. **(a)** Activity snapshot of one example input from SHD in a trained network. Spike raster plots of the input and hidden layer units are shown at the bottom and in the middle. The activity of the twenty readout units is plotted at the top with the brown line corresponding to the correct output for this example. Without any specific regularization, spiking activity in the hidden layer is pathologically high. **(b)** As in (a), but for a network trained with a penalty term for high spiking activity. This form of activity regularization drastically alters the hidden layer activity for the same input, while leaving the winning output of the network unchanged (brown line).

We investigated whether surrogate gradients could instantiate SNNs in this biologically plausible, sparse activity regime. To that end, we trained SNNs with added activity regularization that penalized high spiking activity (Fig. 8b; Methods) and measured the average hidden-layer firing rates and the classification accuracy.

The regularisation term dramatically reduced the number of spikes that were necessary for successful learning. In most cases, the number of spikes could be reduced by approximately two orders of magnitude before there was a notable decline in performance and we found a critical transition in the average number of hidden layer spikes below which networks performed poorly (Fig. 9). For example, in the random manifold task, the transition occurred at ≈ 36 hidden layer spikes per input in a single hidden layer and ≈ 76 spikes in a two hidden layer feed-forward network. In the recurrent network, this number was reduced to 26 (*n*_h_ = 1).

**Fig 9.**
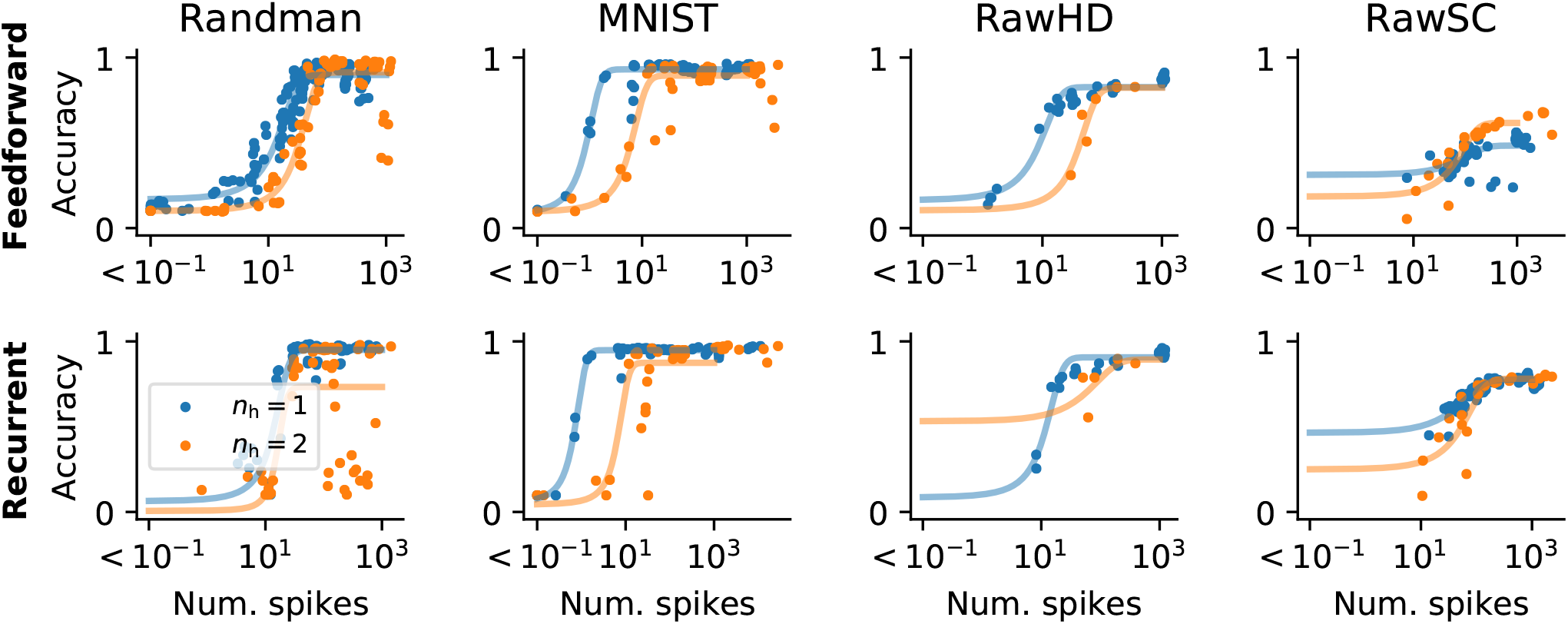
Classification accuracy degrades below a critical number of hidden-layer spikes. Plots showing classification accuracy as a function of the average number of hidden layer spikes per input. The different columns correspond to the different data sets (cf. Fig. 6 and Fig. 7). Top row: Networks with feed-forward connectivity. Bottom row: Networks with recurrent synapses. Blue data points correspond to networks with one hidden layer, whereas orange data points come from networks with two hidden layers. The solid lines correspond to fitted Sigmoid functions.

In the case of MNIST fewer than ten spikes were sufficient on average to achieve the point of diminishing returns beyond which additional spiking activity did not improve classification performance.

This trend could be replicated for all other data sets, with varying degrees of spike reduction. Adding an additional hidden layer generally required more spikes for the same performance. Recurrency generally did not have a great effect on the minimum number of spikes and did not improve performance except on RawSC. On RawSC, 80% classification accuracy was achieved only by a few feed-forward networks with > 2000 spikes on average. In the recurrent network, this level was already attained with ≈ 150 spikes (Fig. 9).

In all cases, the transition from chance level to maximum accuracy occurred in less than one order of magnitude change in the mean number of spikes. On all datasets we tested, the addition of a second hidden layer led to an overall increase in mean spiking activity, which did not yield a notable performance change on Randman and MNIST, but resulted in small improvements on RawHD and RawSC.

These results illustrate that activity regularized SNNs can perform with high accuracy down to some critical activity threshold at which their performance degrades rapidly. Importantly, we found several network configurations that showed competitive performance with an average number of spikes substantially lower than the number of hidden units. For instance, to classify MNIST with high accuracy, an average of 10-20 action potentials was sufficient. Such low activity levels are more consistent with the sparse neuronal activity observed in biological neural circuits and illustrate that surrogate gradients are well-suited to build SNNs which use such lausibly sparse activity levels for information processing.

## Discussion

Surrogate gradients offer a promising way to instill complex function in artificial models of spiking networks. In this article, we focused on two aspects of surrogate gradient learning in SNNs. First, we showed, using a range of supervised classification problems, that surrogate gradient learning in SNNs is robust to different shapes of surrogate derivatives. In contrast, inappropriate choice of scale adversely affected learning performance. Our results imply that for practical applications, surrogate derivatives should be appropriately normalized. Second, by constraining their activity through regularization, we showed that surrogate gradients can produce SNNs capable of efficient information processing with sparse spiking activity.

Surrogate gradients have been used by a number of studies to train SNNs[24], to solve small-scale toy problems with fractionally predictive neurons [43], to train convolutional SNNs on challenging neuromorphic [44, 45] and vision benchmarks [19], or to train recurrent SNNs on temporal problems requiring working memory [21, 22]. These studies used a range of different surrogate derivatives ranging from exponential [21], piece-wise linear [22], or tanh [27], sometimes with a non-standard neuron model with a constant leak term [19], but due to the different function choices and datasets they are not easily comparable. Here we provide such a comprehensive comparison.

Towards this end, we had to make some compromises. Like previous studies, our study is limited to supervised classification problems, because supervised learning offers a well-defined and intuitive quantification of computational performance. To keep the number of model parameters tractable, we focused on current-based LIF neurons. Moreover, we entirely dispensed with Dale’s law and relied solely on all-to-all connectivity. However, we believe that most of our findings will carry over to more realistic neuronal, synaptic, and connectivity models. The present study thus provides a set of blueprints and benchmarks to accelerate the design of future studies.

Although considered biologically implausible [46], we have limited our study to training SNNs with backpropagation, the de-facto standard for computing gradients in systems involving recurrence and hidden neurons. Although there exist more plausible forward-in-time algorithms like real-time recurrent learning (RTRL) [47] they are prohibitively expensive or else require additional approximations that affect learning performance [24, 25, 48, 49]. Instead, using BPTT enabled the targeted manipulation of gradient flow through different elements of the network which allowed us to dis-entwine some of complexity of surrogate gradient learning. Finally, the present study was purely numerical. It will require future work to establish a rigorous theoretical understanding of surrogate gradient learning dynamics to understand optimal derivatives and algorithms for learning in SNNs.

In summary, surrogate gradients allow translating the success of deep learning to biologically inspired SNNs by optimizing their connectivity toward functional complexity through end-to-end optimization. The in-depth study of both surrogate gradients and the resulting functional SNNs will occupy scientists for years to come and will likely prove transformative for neural circuit modeling.

## Methods

### Supervised learning tasks

In this study we used a number of synthetic and real-world learning tasks with the overarching aim to balance computational feasibility and practical relevance. To that end, we focused on synthetic datasets generated from random manifolds and real-world auditory datasets.

### Smooth random manifold datasets

We generated a range of synthetic classification data sets based on smooth random manifolds. Suppose we want to generate a smooth random manifold of dimension *D* in an embedding space of dimension *M*. We are looking for a smooth random function *f*: ℝ^*D*^ → ℝ^*M*^ defined over the finite interval 0 ≤ *x* < 1 in each of intrinsic manifold coordinate axis. Moreover, we would like to keep its values bounded in a similar (0 ≤ *x* < *τ*_randman_)^*M*^ box in the embedding space. To achieve this we first generate *M* smooth random functions *f_i_*: ℝ^*D*^ → ℝ and then combine them to *f*: ℝ^*D*^ → ℝ^*M*^. Specifically, we generate the *f_i_* from the Fourier basis as follows:

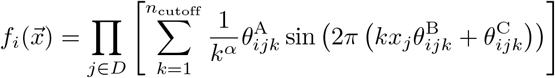

where the parameters 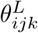 for *L* ∈ {A, B, C} were drawn i.i.d. from a uniform distribution 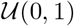. We set *n*_cutoff_ = 1000 which leaves *α* as a parameter that controls the smoothness of the manifold. Larger values of *α* lead to more slowly varying manifolds whereas smaller values increase the high frequency content (Fig. 1a). In addition to *α*, the complexity of the learning problem can be seamlessly adjusted by either increasing the intrinsic dimension *D* of the manifold or the number of random manifolds whereby each random manifold corresponds to a separate class of the classification problem (Fig. 1b). We generated spike trains by randomly sampling the resulting random manifolds and standardizing their numerical values to lie between 0 and *τ*_randman_ in all embedding dimensions. Finally, the resulting values were interpreted as the firing times from one class of the classification problem (Fig. 1c).

We chose a default parameter set for most of our experiments that struck a good balance between minimizing both the embedding dimension and the number of samples needed to solve the problem while simultaneously not being solvable by a two layer network without hidden units. Specifically, we chose a 10-way problem with *D* = *α* = 1, *M* = 20, and *τ*_randman_ = 50ms and fixed the random seed in situations in which we reused the same dataset. We generated 1000 data points for each class out of which we used 800 for training and two sets of 100 each for validation and testing purposes.

### Spike latency MNIST dataset

To convert the analog valued MNIST dataset [50] to firing times we proceeded as follows. We first standardized all pixel values 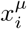 to lie within the interval 0 ≤ *x* < *τ*_eff_. We then computed the time to first spike latency *T* as the time to reach firing threshold of a leaky integrator

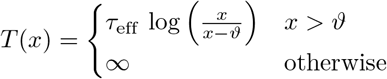

in our simulations we used *ϑ* = 0.2 and *τ*_eff_ = 50ms (Fig. 7a,b).

### Auditory datasets

We used both spiking and non-spiking auditory datasets of digit and word utterances. Specifically, we used the SHDs without any further preprocessing [23]. For performance reasons and to dispense with the spike conversion process, we ran additional simulations with non-spiking auditory inputs. Specifically, we worked with the raw Heidelberg Digits (RawHD) and Pete Warden’s Speech Commands dataset (RawSC) [36] which were preprocessed as follows: We first applied a pre-emphasis filter to the raw audio signal *x*(*t*) by computing *y*(*t*) = *x*(*t*) − 0.95*x*(*t* − 1). We then computed 25ms frames with a 10ms stride from the resulting signal and applied a Hamming window to each frame. For each frame we computed the 512-point fast Fourier transform to obtain its power spectrum. From the power spectrum we further computed the filter banks by applying 40 triangular filters on a Mel-scale [51]. After cropping or padding to 80 (RawHD) or 100 (RawSC) steps by repeating the last frame, the analog valued filter banks were fed directly to the SNNs.

### Network models

To train SNN models with surrogate gradients we implemented them in PyTorch [33]. To that end, all models were explicitly formulated in discrete time with time step Δ*t*.

### Neuron model

We used leaky integrate-and-fire neurons with current-based exponential synapses [52, 53]. The membrane dynamics of neuron *i* in layer *l* were characterized by the following update equations

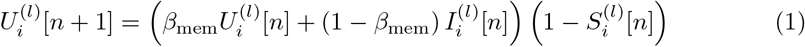

where 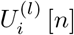 corresponds to the membrane potential of neuron *i* in layer *l* at time step *n* and is its 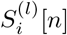 the associated output spike train defined via the Heaviside step function Θ as 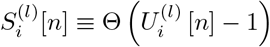. Note that in this formulation, the membrane dynamics are effectively re-scaled such that the resting potential corresponds to 0 and the firing threshold of 1. This choice simplifies the implementation of the neuronal reset dynamics through the factor on the right hand side. During the backward pass of gradient computation with BPTT, the derivative of the step function is approximated using a surrogate as explained later. In situations in which we ignored the reset term this was done differently for the output spike train and the spike train underlying the reset term, by detaching it from the computational graph. The membrane decay variable *β*_mem_ is associated with the membrane time constant *τ*_mem_ through 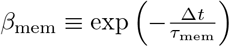. Finally, the variable 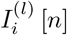 is the synaptic current defined as follows

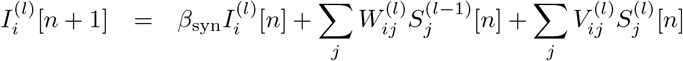

with feed-forward afferent weights *W_ij_* and the optional recurrent weights *V_ij_*. In analogy to the membrane decay constant, *β*_syn_ is defined as 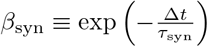. We set *τ*_mem_ = 10ms and *τ*_syn_ = 5ms. Together the computations involved in each time step can be summarized in the computational graph of the model (cf. Fig. 4).

### Readout layer

The readout units in our models are identical to the above neuron model, but without the spike and associated reset. Additionally, we allow for a separate membrane time constant *τ*_readout_ = 20ms with 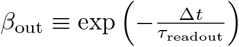. Overall their dynamics are described by

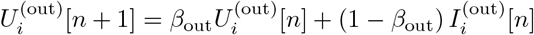

### Connectivity and initialization

We used all-to-all connectivity in all simulations without bias terms unless mentioned explicitly. The weights were initialized from a uniform distribution 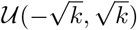 with 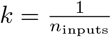 where *n*_inputs_ is the number of afferent connections.

### Readout heads and supervised loss function

We trained all our networks by minimizing a standard cross entropy loss 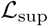 defined as

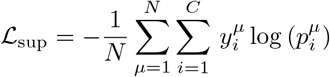

where 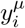 is the one-hot encoded target for input *μ*, *N* is the number of input samples, and *C* is the number of classes. The output probabilities 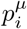 were given by the Softmax function

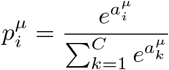

in which the logits 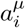 for each input *μ* were given by either defined as the maximum 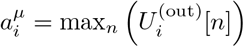, a notion inspired by the Tempotron [29], or the sum over all time steps 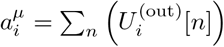 depending on the readout configuration we used.

### Activity regularization

To control spiking activity levels in the hidden layers we employed two forms of activity regularization. First, to prevent quiescent units in the hidden layers, we introduced a lower activity threshold *ν*_lower_ at the neuronal level defined as

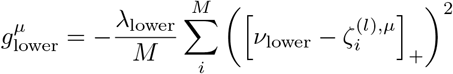

with the neuronal spike count 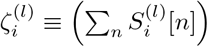 and the number of neurons *M* in hidden layer *l*. Similarly, we defined an upper threshold at the population level as

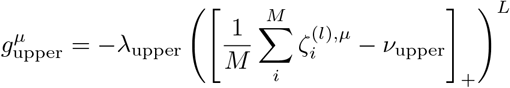

for which we explored both values of *L* ∈ {1, 2}. The overall regula)rization loss was computed by summing and averaging 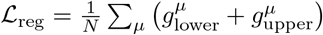. Finally we optimized the total loss 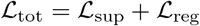 by dint of surrogate gradient descent.

### Surrogate gradient descent

We minimized the loss 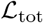 by adjusting the parameters *W* and *V* in direction of the negative surrogate gradients using Adam with default parameters [54]. Surrogate gradients were computed with back-propagation through time using PyTorch’s automatic differentiation capabilities. To deal with the non-differential spiking nonlinearity of the hidden layer neurons, we approximated their derivatives 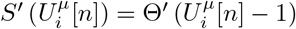 with suitable surrogates 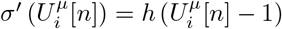. Throughout, we used the following functions *h*(*x*):

**SuperSpike:** 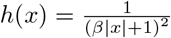
**Sigmoid′:***h*(*x*) = *s*(*x*) (1 − *s*(*x*)) with the Sigmoid function 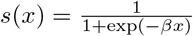
**Esser et al.:** *h*(*x*) = max(0, 1.0 − *β*|*x*|)

where *β* is a parameter that controls the slope of the surrogate derivative. In Figure 5, we additionally considered an asymptotic variant of SuperSpike defined as 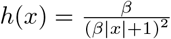. Unless mentioned otherwise we set *β* = 10. Table 1 specifies the relevant hyperparameters used in the different simulations. An example of how these manipulations can be achieved easily with PyTorch can be found at https://github.com/fzenke/spytorch [55]. All simulations and parameter sweeps were performed on compute nodes equipped with Nvidia Quadro RTX 5000 and V100 GPUs.

**Table 1.** Parameter values of network simulations.

| Parameter | Randman | MNIST | SHD | RawHD | RawSC |
| --- | --- | --- | --- | --- | --- |
| Number of input units | 20 | 784 | 700 | 40 | 40 |
| Number of hidden units | 100 | 100 | 256 | 256 | 256 |
| Number of readout units | 10 | 10 | 20 | 20 | 35 |
| $\Delta t$ / Number of steps | 1ms / 100 | 1ms / 100 | 2ms / 500 | 2ms / 80 | 2ms / 100 |
| Minibatch size | 128, 250 | 256 | 256 | 128 | 512 |
| Number of epochs | 50–100 | 100 | 200 | 50 | 50 |
| Dataset (train/valid./test) | 8k/2k/2k | 45k/5k/10k | 7498/833/2088 | 7498/833/2088 | 84849/9981/11005 |
| Number of hidden layers $n_h$ | $\leq 2$ | $\leq 3$ | $\leq 2$ | 1 | 1 |
| Learning rate sweep | $10^{-3} \leq \eta \leq 0.1$ | $2 \times 10^{-4} \leq \eta \leq 1^{-1}$ | $2 \times 10^{-4} \leq \eta \leq 1^{-1}$ | $1 \times 10^{-3}$ | $1 \times 10^{-3}, 5 \times 10^{-3}, 0.01$ |
| Best $\eta$ | 0.05 | 0.01 | $1 \times 10^{-3}$ | $1 \times 10^{-3}$ | $1 \times 10^{-3}$ |
| Lower L2: $\lambda_{\text{lower}} / \nu_{\text{lower}}$ | $100.0/10^{-3}$ | — | $100.0/10^{-3}$ | $100.0/10^{-3}$ | $100.0/(0.01, 10^{-3})$ |
| Upper L1 strength $\lambda_{\text{upper},1}$ | 1,100 | 0–100 | 0.06 | 0, 10, 20, 100 | 0.06, 1, 10 |
| Upper L1 threshold $\nu_{\text{upper},1}$ | 0–1000 | 0–1000 | 0–1000 | 0, 0.1, 0.2, 0.5, 1, 100 | 0–1000 |
| Upper L2 strength $\lambda_{\text{upper},2}$ | 0,1,100 | 0, 0.001, 0.1, 1 | — | — | 0,1 |
| Upper L2 threshold $\nu_{\text{upper},2}$ | 0–100 | 0.1, 100 | — | — | 10,100 |

## Acknowledgments

FZ was supported by the Wellcome Trust (110124/Z/15/Z) and the Novartis Research Foundation. TPV was supported by a Wellcome Trust Sir Henry Dale Research fellowship (WT100000), a Wellcome Trust Senior Research Fellowship (214316/Z/18/Z) and an ERC Consolidator Grant SYNAPSEEK.

